# *PointedSDMs* – an R package to help facilitate the construction of integrated species distribution models

**DOI:** 10.1101/2022.10.06.511075

**Authors:** Philip S. Mostert, Robert B. O’Hara

## Abstract

1. Integration of disparate datasets is a rapidly growing area in quantitative ecology, and is subsequently becoming a major asset in understanding the shifts and trends in species’ distributions.
2. However, the tools and software available to construct statistical models to integrate these disparate datasets into a unified framework is lacking, stagnating the growth of data integration in applications.
3. This paper presents *PointedSDMs*: an easy to use *R* package used to construct integrated species distribution models using the integrated nested Laplace Approximation methodology.
4. The package is presented in this paper through an illustrative example for a selection of species found across Pennsylvania state.

## 2 Introduction

Ecological research in the 21^st^ century has been characterized by the mass accumulation of species occurrence data worldwide, due mainly to the incessant advancements in digital technology and online data repositories, whose aim is to increase the availability of biodiversity data through the internet and other accessible data sharing protocols (LaDeau et al., 2017). While this accumulation of data has certainly expanded the potentials of ecological analysis on the spread, range shifts and relationship species have with the underlying environment, a multitude of challenges have arisen as a result: the data are likely to have come from disparate sources, resulting in heterogeneous attributes, assumptions and sampling protocols inherent in each individual dataset (Fletcher Jr et al., 2019).

As a result, a myriad of statistical methods to perform analysis, make predictions and efficiently use all available data have been produced – so called integrated species distribution models (ISDMs) in literature (see Miller et al., 2019, for a detailed review).

Although the fundamental methodology and implementation between the developed methods to combine data may differ, a common result is that integrated data models appear to not only expand the spatial scope of a study, but appear to be superior to models with a single data source by providing improvements to the results and estimates in comparison to using only a single dataset (see for example: Fithian et al., 2015; Bowler et al., 2019; Miller et al., 2019).

Although IDMs are being developed, a significant problem in the uptake of integrated models however is the lack of general software and tools to make inference with them – and as a result, the overall uptake of data integration across the fields of ecology has been generally slow.

Here we introduce *PointedSDMs*, an easy to use *R* (R Core Team, 2022) package designed to fit complex marked SDMs using data obtained from heterogeneous sources, and integrate them all together in a unified statistical framework. It does so using a hierarchical state space formulation – in which we link a process model (which provides a description of the true distribution of the model) with observation models for each dataset, dependent on their underlying sampling protocols (Isaac et al., 2020).

The integrated model for this package is fitted using integrated nested Laplace approximation (INLA) – a computationally efficient method used by Bayesian statisticians to fit latent Gaussian models. The theory behind the INLA methodology is discussed in detail in (Rue et al., 2009), which may appear intimidating for researchers with a limited background in mathematics. However fitting models with this methodology is made simple in an accessible interface with the now established *R-INLA* package (Martins et al., 2013). The *PointedSDMs* package constructs a wrapper around the R package *inlabru* (Bachl et al., 2019), which in turn builds off the *R-INLA* package to help provide a user-friendly method to simplify the modeling of spatial process models.

### 2.1 Statistical model

The aim of our state-space point process model is to use the available species’ location data to make inference about the “true” distribution of the population of the species’ (Isaac et al., 2020). To do so, we use a hierarchical modeling structure with an underlying process model which describes how points are distributed in space. This process has an intensity function (denoted here by λ(*s*)) which is some function of environmental covariates ***X*** and parameters ***θ*** such that a higher intensity implies that the species is more abundant in a location.

For this model, we assume that the underlying process model is a log-Gaussian Cox process (LGCP) with an intensity function given as λ(*s*) = *exp*{*η*(*s*)}, which describes the expected number of species at some location, *s*. The log of this intensity function is thus given as:

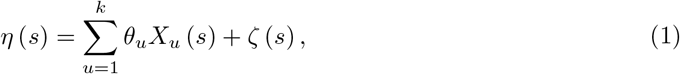

where *ζ*(*s*) is a zero-mean spatially continuous Gaussian random field, included in the model to account for potential spatial autocorrelation and the effects of all the environmental covariates not included in the model. Therefore, the expected number of species’ presences within a region Ω is given by the integral of the intensity function across the entire region:

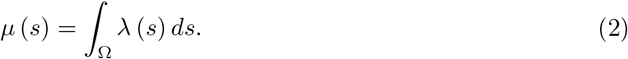

Next, we assume that each dataset process (*Y_i_*, *i* = 1, 2, …, *n*) has its own sub-model (observation model), which links the intensity function to the dataset’s assumed likelihood, given by, 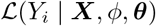, where *ϕ* are parameters assumed for the underlying process model.

Isaac et al. (2020) presents a succinct overview of three commonly used data types typically found in point-process SDMs: abundance data (modelled as a Poisson random) variable, presence only data (modelled as a thinned Poisson random variable) and presence absence data (modelled as a Bernoulli random variable with a *cloglog* link function (see Kéry & Royle, 2016). In addition to the species location data, datasets sometimes include additional trait variables (often referred to as marks). These data may also be included in the point-process modelling framework to supplement the amount of information in the SDM.

Then, by combining the process model with the observation models, the full likelihood for the data processes ***Y*** = {*Y*_1_, *Y*_2_, …, *Y_n_*} is given by:

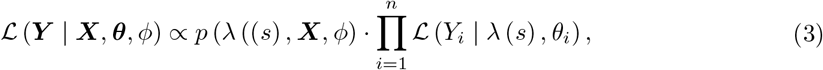

that is, the model component for the latent state of the model, multiplied by the product of the individual likelihoods for the data processes.

## 3 Package functionality

*PointedSDMs* was developed to streamline the modelling process and provide a general framework for integrated species distribution models for ecologists who have a collection of heterogeneous datasets at hand. It does so by re-formatting and assigning appropriate metadata to the species’ location and covariate data, and then constructing the relevant objects required by *R-INLA* (Martins et al., 2013) to do the model fitting. The package contains four primary functions for model pre-preparations (`intModel`), fitting and inference (`fitISDM`) and cross-validation (`datasetOut` and `blockedCV`), as well as several generic functions related to plotting, printing and predicting the results of the model.

`intModel` is the first function used in the integrated modeling process, and is built using *R*’s *R6* (Chang, 2021) object orientated system. Here, the user adds the species location data, environmental covariates, as well as additional *R-INLA* and *sp* (Pebesma & Bivand, 2005) objects required; most of the other arguments for this function are used to define variable names and terms to be included in the model. Since this is an *R6* object, there are a handful of slot functions which allow further specification and adjustments of the components in the model. *PointedSDMs* allows datasets from three sampling schemes: presence only, presence absence and counts data, where the latter two are defined in the model through their response variable names, using the `intModel`’s arguments *responsePA* and *responseCounts* respectively.

If the user defines a spatial partitioning of their data points using `intModel`’s slot function, `‘.$spatialBlock‘`, spatial cross-validation may be performed using the function, `blockedCV`: which iteratively calculates a cross-validation score by leaving a certain block of data out of the model based on their spatial location.

`fitISDM` is used for the modeling and estimation of the integrated model. The *data* argument of the function is an object created by the function `intModel`, which contains the necessary information and metadata required in the model. The second argument, *options* is used to control any additional *R-INLA* or *inlabru* options.

After the model has been estimated, another form of spatial cross-validation may be completed using the function, `datasetOut`. The function works by calculating a cross-validation score from the following steps:

1. Running a new model with one less dataset (from the main model) – resulting in a reduced model,
2. predicting the intensity function at the locations of the left-out dataset with the reduced model,
3. using the predicted values as an offset in a new model,
4. finding the difference between the marginal-likelihood of the main model (i.e the model with all the datasets considered) and the marginal-likelihood of the offset model.

Installation of the package may be done directly from *CRAN* servers using the following *R* script:

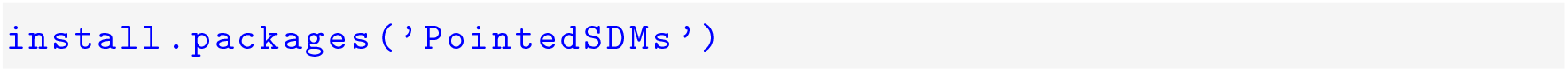

Any concerns and questions regarding the use of the package may be asked on issues board of the package’s GitHub repository: https://github.com/PhilipMostert/PointedSDMs.

## 4 Worked example

### 4.1 Introduction

The example below illustrates the use of *PointedSDMs* in a worked example, using datasets from a variety of distinct sources. The datasets used include observations of *Setophaga* (genus) from both structured and unstructured sampling schemes, obtained from various locations around Pennsylvania state (41°12’N, 77°11’W) on the eastern side of the United States of America (USA), which were collected between 2005 and 2009. Similar studies using these data were presented by Miller et al. (2019) and Isaac et al. (2020), who used the *WinBUGS* (Lunn et al., 2000) and the *R-INLA* package respectively to obtain results.

### 4.2 Description of datasets

The data used in this analysis comes from three heterogeneous sources, where we assumed the underlying sampling protocol for each dataset is unique to that dataset (we considered datasets representing: presence only, presence absence and counts frameworks).

The citizen science PO data were obtained from *eBird* (Sullivan et al., 2009), a citizen science project launched by the *Cornell lab of Ornithology* where amateur birders are able to submit checklists of avian detections to an online data repository, which has grown significantly since its inception, and has established itself as a significant tool in scientific research. Since the *eBird* data is collected by non-scientists, as so we expect the biases typically found in CS data.

The other two datasets used come from structured survey data. The first comes from the *North American Breeding Bird Survey* (BBS) (Pardieck & Hudson, 2018), a long-term birding project designed to monitor changes in North American breeding bird populations for numerous species (Sauer et al., 2017). For their analysis, Isaac et al. (2020) treated the BBS data as a replicate presence-absence data per sight, however for illustrative purposes we treat it as a count datasets, with a response variable denoting the number of species observed at each sight.

The *Pennsylvania Breeding Bird Atlas* (BBA) (obtained from the supporting information file in (Miller et al., 2019)) is another long-term avian project following a standardized collection process (Wilson et al., 2012). Following Isaac et al. (2020), we treat this data as PA and assume that the sights are small enough to be represented as points.

Two standardized and continuous spatial covariates describing the study area were used in this analysis. The first, elevation, describes the height in meters above sea level, obtained from the package, *elevatr* (Hollister et al., 2021), and the second, canopy, describes the percentage of tree canopy covered in the area, obtained from the package, *FedData* (Bocinsky, 2022), which in turn accesses the data from the *National Land Cover Database* (NLCD). Both of these packages produced spatial covariates in the form of *Raster* objects, which we stacked into a single *RasterBrick* object before analysis.

**Figure 1:**
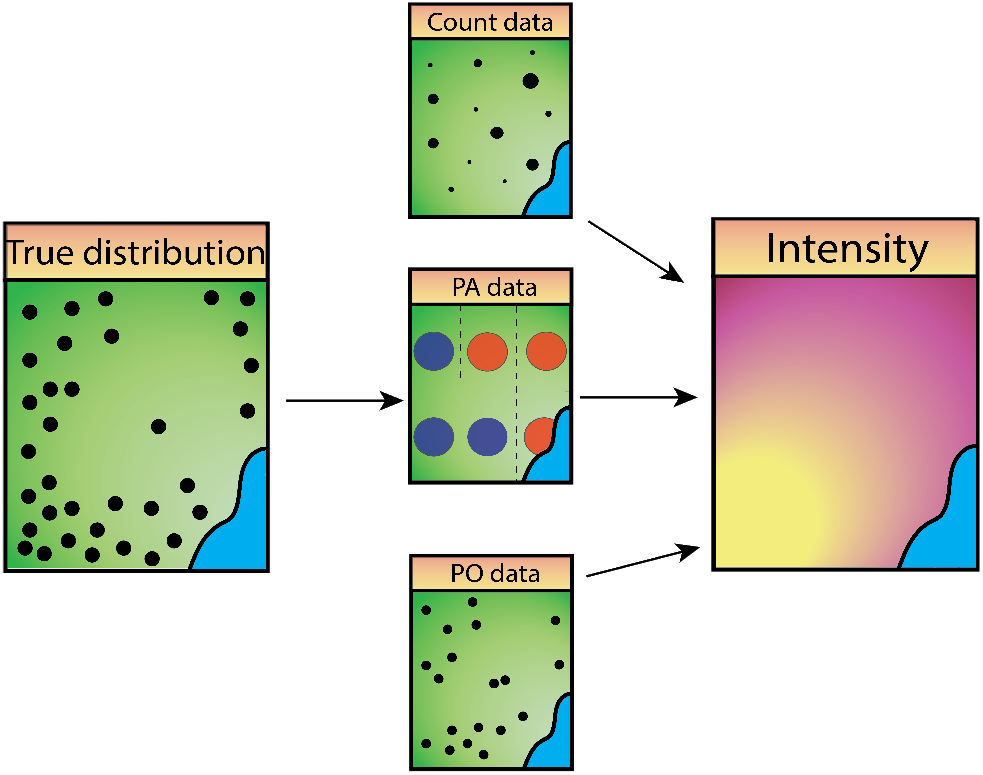
Representation of the structure of the integrated species distribution model, where each dataset is a separate realization of the “true” species distribution. This is done by assuming each dataset has its own observation process, with a common latent, which is described by ecological covariates and parameters.

### 4.3 Model preparations

The first step to running an integrated model with *PointedSDMs* is to organize and assign appropriate metadata to the individual datasets, using the `intModel` function, which is used to initiate and prepare the statistical model before any inference is made; and so the arguments it takes are used to assign the relevant metadata to the datasets and covariates as well as set up all the objects required by *R-INLA*.

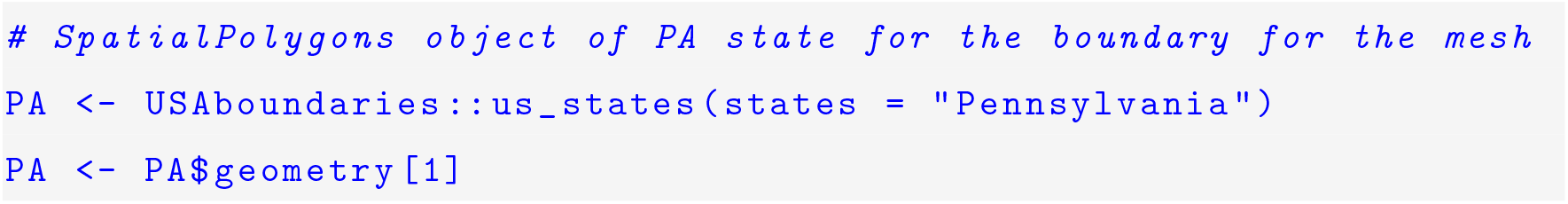
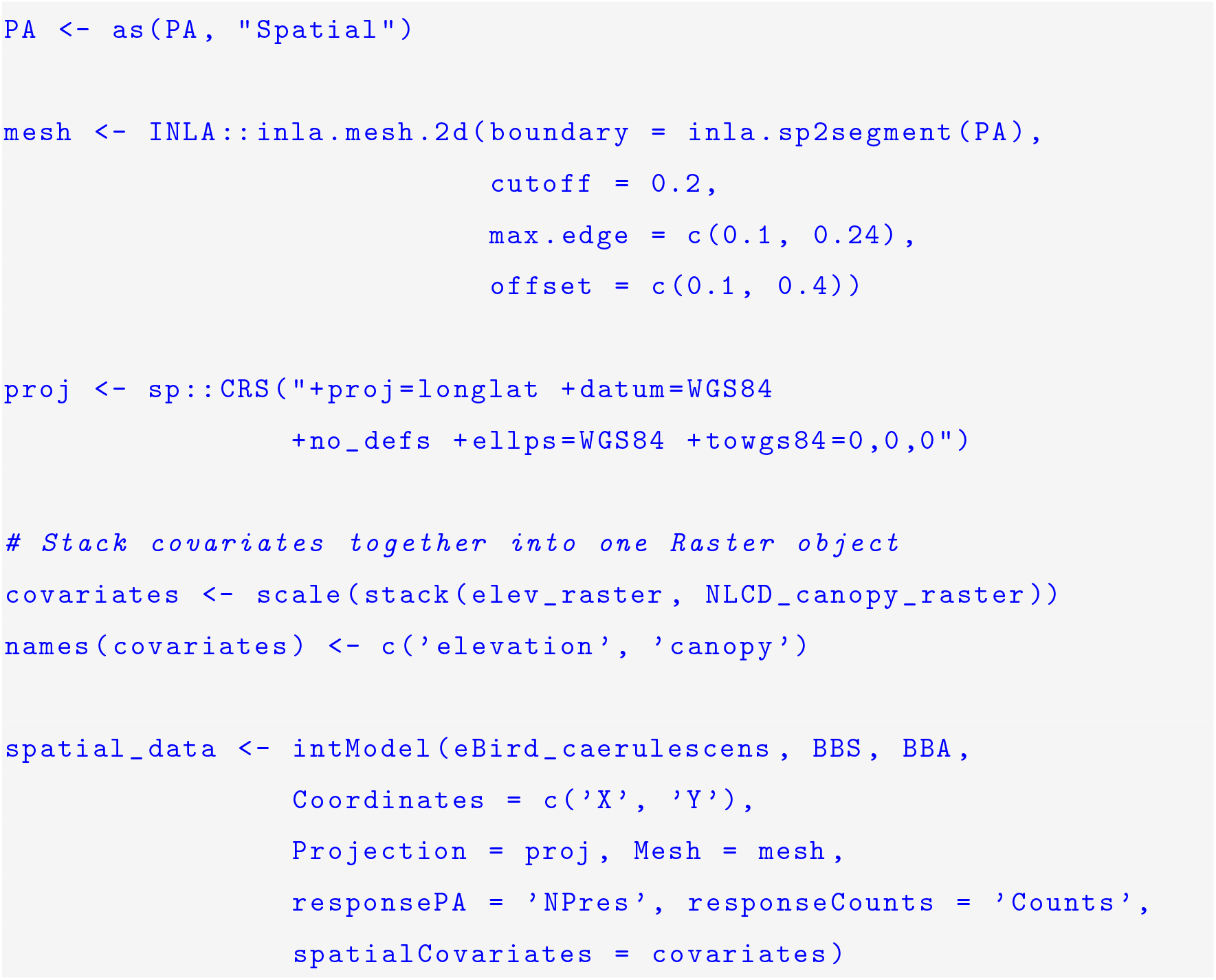

We would also like to approximate spatial autocorrelation in the model not accounted for by our covariates, which may be computationally expensive for large point process models. To counter this issue, *R-INLA* approximates the continuous spatial field with a triangulated mesh. The mesh for this example was created with the `inla.mesh.2d` function by supplying a *SpatialPolygons* boundary as well as the *max.edge, offset*, and *cutoff* arguments.

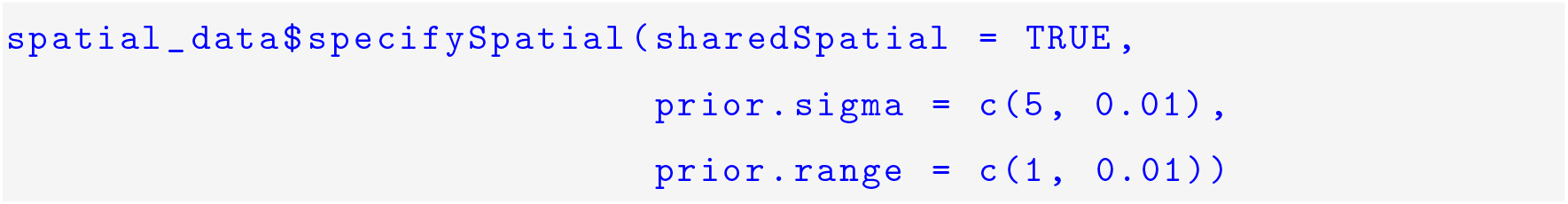

The stochastic partial differential equation (SPDE) models were specified using penalizing complexity (PC) priors (Simpson et al., 2017), which are designed to control the spatial range and standard deviation in the GRF’s Matérn covariance function in order to reduce over-fitting in the model.

Simmonds et al. (2020) demonstrated in a simulation study that running a second spatial field for opportunistically collected PO data is a useful method to account for bias when knowledge of the sources of bias is scant or when covariates to adjust for bias are unavailable. Therefore, we use the `‘.$addBias‘` function to add a second spatial field to our CS data to account for potential biases not reflected in the shared field.

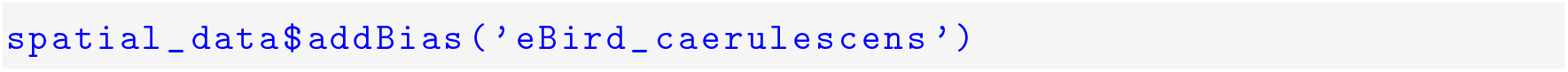

### 4.4 Results

The integrated model is easily fit using the `fitISDM` function as below, which takes two arguments: data (which is an *intModel* object created above), and options (which is a list of *R-INLA* and *inlabru* options used to configure the model). In this model, the two fixed covariates and separate intercept terms for the three datasets were considered. In addition to the bias field for *eBird_caerulescens*, a common spatial field was used across the datasets; and to speed up computation time, *R-INLA’s* empirical Bayes’ integration strategy was used.

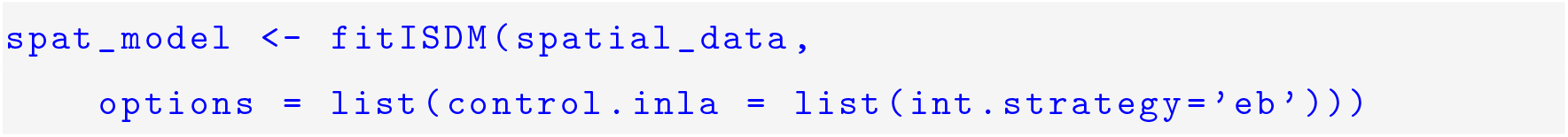

*PointedSDMs* also includes the function `datasetOut`, which iteratively omits one dataset (and its associated marks) out of the full model. Table 2 below illustrates the results of omitting one dataset out of the model at a time, where the mean change in fixed effects appears to vary significantly between the datasets.

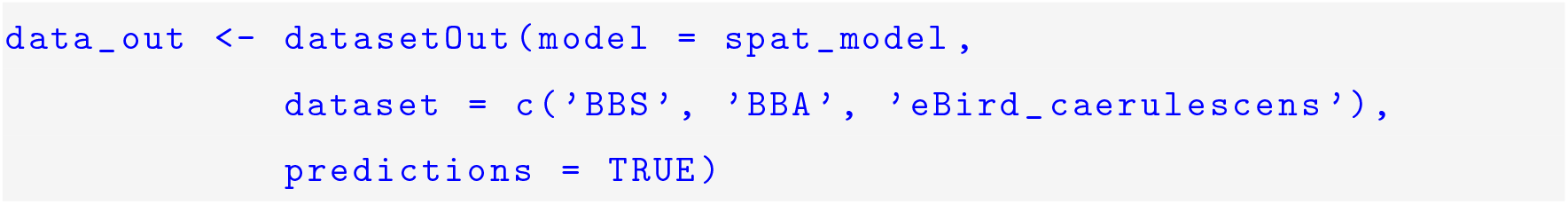

**Table 1:** Table showing the summary of the fixed effects from the integrated model for species caerulescens.

| Fixed effects | Mean | SD | 0.025quant | 0.5quant | 0.975quant |
| --- | --- | --- | --- | --- | --- |
| *Elevation* | 1.31 | 0.09 | 1.13 | 1.31 | 1.49 |
| *Canopy* | 0.06 | 0.06 | -0.05 | 0.06 | 0.17 |
| *eBird intercept* | -0.60 | 1.38 | -3.30 | -0.60 | 0.17 |
| *BBS intercept* | 0.48 | 0.50 | -0.50 | 0.48 | 1.45 |
| *BBA intercept* | -5.70 | 0.51 | -6.71 | -5.70 | -4.70 |

**Table 2:** Table showing the leave one out cross-validation score as well as the changes in fixed effects as a result of leaving a dataset out. The cross-validation score is calculated by finding the difference between the marginal likelihood of the full model, and the marginal likelihood of the model with the dataset left out.

| Dataset left out | ΔElevation | ΔCanopy | Cross-validation score |
| --- | --- | --- | --- |
| BBS | 0.1974 | 0.1074 | 3.4660 |
| BBA | -0.5537 | -0.1786 | 2.7370 |
| eBird | 0.0951 | 0.0867 | 2.3187 |

Setting `predictions = TRUE` allows the user to calculate a cross-validation score obtained by leaving out a dataset. In this case leaving out the BBS dataset causes the greatest difference in marginal likelihood between the main model and the reduced (without BBS) model, suggesting that this dataset provides the most information in our integrated model.

### 4.5 Predictions

A crucial part of the process of making SDMs is creating prediction maps to help researchers understand the species’ spread. Predictions of the ISDMs from `fitISDM` are made easy using the `predict` function. The function will automatically create individual formulas to predict per dataset after the user has specified which components they would like to predict (with the arguments: *covariates, spatial*, and *intercept*); and all components used in the model may be included by setting the *predictor* argument to `TRUE`. However any formula may be predicted by using the function’s *formula* argument.

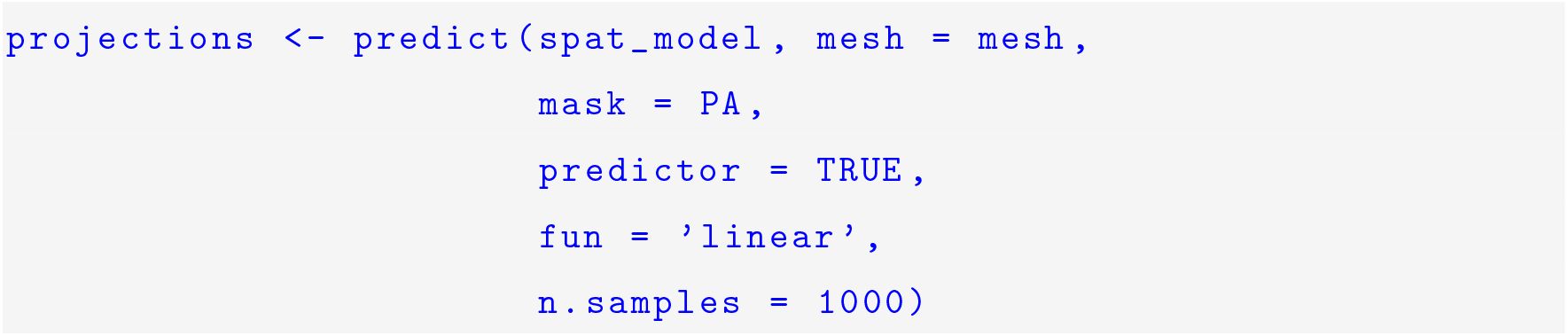

**Figure 2:**
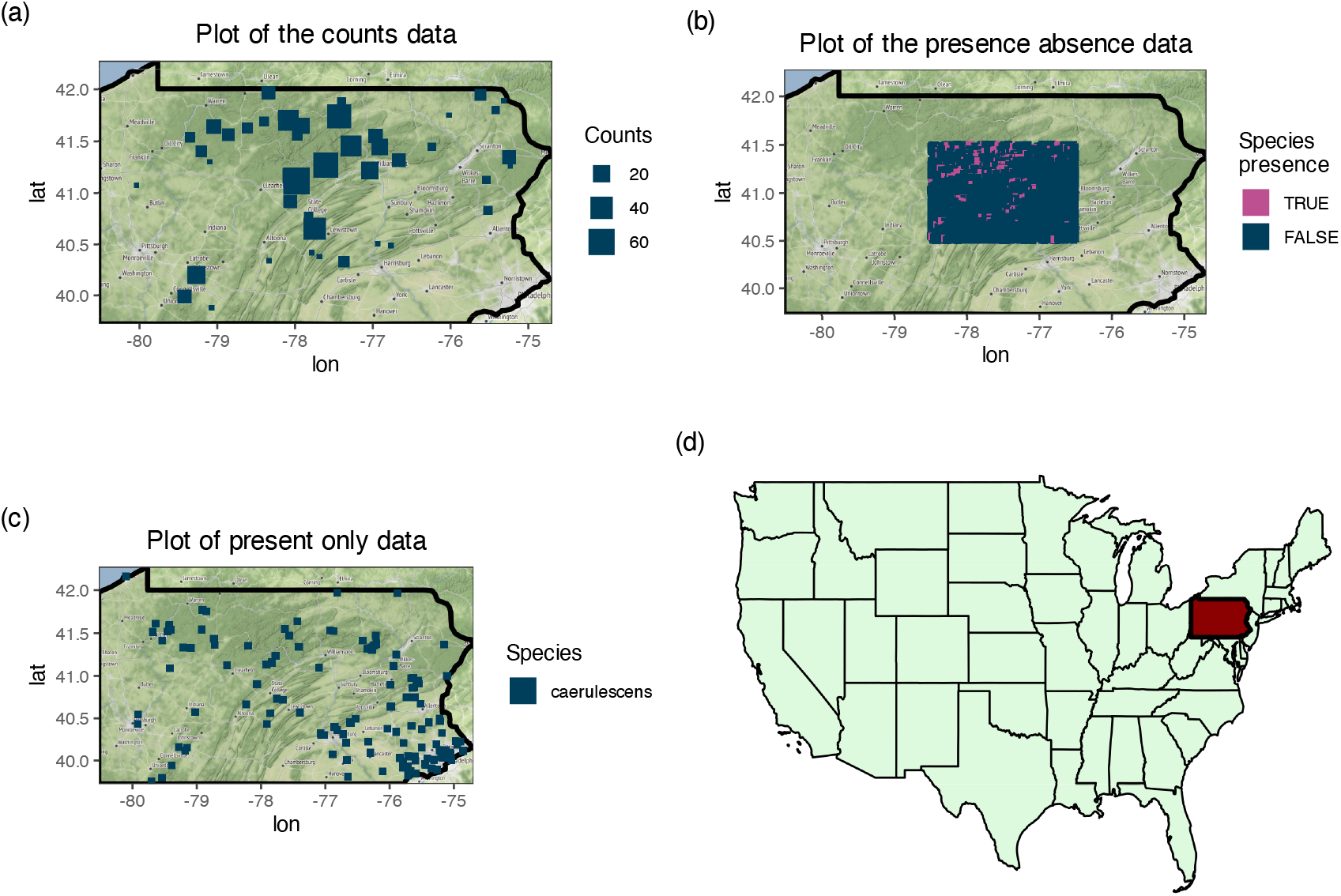
(a)-(c): Plots of the datasets based on their underlying sampling protocol. (d) Map of the United States of America, highlighting Pennsylvania state.

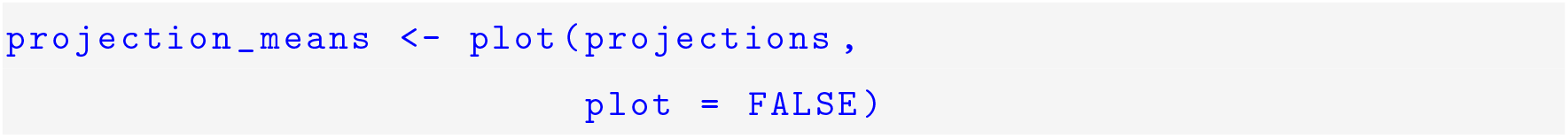

*PointedSDMs* also provides methods to plot basic predictive maps for a variety of statistics. By setting the *plot* argument to `FALSE`, the *ggplot* (Wickham, 2016) object of the predicted statistic is given, which would allow for more custom plotting functionality.

## 5 Conclusions

*PointedSDMs* is an *R* package that provides the tools to make the most of the vast volume of species location data available today, by promoting and facilitating the integrated modeling of marked point process SDMs in a convenient way. It does so using the now well-established INLA methodology, and by constructing wrapper functions around the R package, *inlabru*.

## Opportunities for future work

Different groups of species influence each other through a multitude of processes (such as predation and competition), thereby affecting each others distribution across space and time. A method to account for these processes would be to add inter-species interactions between different species, therefore changing the model framework to a joint species distribution model (JSDM).

A limitation of this model is that it only incorporates a small subset of the types of data used in ecology (presence only, presence absence and count data). Therefore, there is an opportunity to incorporate other types of data (such as biomass and movement data) into this framework, which would thereby extend the possibilities of research within a project.

Furthermore, allowing the user to add their own custom non-linear and smoothing components to the model is essential, given that complex ecological studies often require these terms.

Finally, constructing the necessary tools and data pipelines to move species and environmental data from online repositories to create a complete workflow in a way that is not only reproducible, but also easy enough to use for ecologists and policy makers with minimal basic skills, would allow a package like this to show off its full potential.

## Authors’ contributions

P.S.M wrote the script for the R package, *PointedSDMs*, provided the graphics and led the writing for the first draft of the manuscript. R.B.O provided conceptualization of the project, supervision and reviewed the manuscript.

## Data availability statement

All data are freely available for the reader in the *R* package, *PointedSDMs*, which is available on the Comprehensive R Archive Network: https://cran.r-project.org/web/packages/PointedSDMs/index.html, the *R* code is also open source and available on GitHub: https://github.com/PhilipMostert/PointedSDMs. The version of the package used for this manuscript (v1.1.1) is archived on Zenodo (Mostert and O’Hara, 2022, 10.5281/zenodo.7078847).

## Acknowledgements

We want to thank Walter Jetz and Petr Keil for help and discussions early in this project.

## Conflict of interest

We declare no conflicts of interest.

## Notes

https://github.com/PhilipMostert/PointedSDMs

### Competing Interest Statement

The authors have declared no competing interest.

